# The involvement of HDAC3-mediated inhibition of microRNA cluster 17-92 in hyperoxia-mediated impairment of lung development in neonatal rats

**DOI:** 10.1101/2020.03.01.971663

**Authors:** Di Wang, Hui Hong, Xiao-Xia Li, Jing Li, Zhi-Qun Zhang

**Affiliations:** Department of Pediatrics, Affiliated Hangzhou First People’s Hospital, Zhejiang University School of Medicine, Hangzhou 310006, P. R. China; Department of Neonatology, Affiliated Hangzhou First People’s Hospital, Zhejiang University School of Medicine, Hangzhou 310006, P. R. China

**Keywords:** Histone deacetylase 3, microRNA-17-92 cluster, Placental growth factor, Bronchopulmonary dysplasia, Pulmonary angiogenesis, Alveolarization

## Abstract

**Background:** The incidence of bronchopulmonary dysplasia (BPD), a chronic lung disease of newborns, has been paradoxically rising despite medical advances. Histone deacetylase 3 (HDAC3) has been reported to be a crucial regulator in alveologenesis. Hence, this study aims to investigate the mechanism of HDAC3 in the pulmonary angiogenesis and alveolarization of BPD.

**Methods:** A hyperoxia-induced mouse model of BPD was constructed. The mean liner intercept (MLI) and alveolar volume were measured to evaluate the alveolarization in BPD mice. Immunofluorescence assay was performed to detect the microvessel density (MVD) of lung tissues. Next, the expression of HDAC3 and its enrichment in the promoter region of microRNA (miR)-17-92 cluster, as well as the enrichment of p65 in the placental growth factor (Pgf) promoter region were detected by Western blot analysis and chromatin immunoprecipitation (ChIP) assay. The effect of HDAC3 and p65 on the activity of miR-17-92 promoter and Pgf promoter were examined by dual-luciferase reporter gene assay, respectively. Finally, the role of HDAC3 in angiogenesis and alveolarization through miR-17 regulated EZH1-p65-Pgf axis was validated in BPD mouse models.

**Results:** HDAC3 was involved in the regulation of alveolarization and angiogenesis in BPD. Results demonstrated that the expression of the miR-17-92 cluster in BPD was regulated by HDAC3. miR-17 was related to the regulatory role of HDAC3 in regulating EZH1 expression and in lung fibroblasts of BPD. Besides, results showed that EZH1 could promote Pgf expression by recruiting p65 to regulate BPD. HDAC3 regulated the expression of EZH1 through miR-17 to promote the recruitment of p65 in the Pgf promoter region, thus enhancing the transcription and expression of Pgf. HDAC3 was demonstrated to regulate Pgf through the miR-17-EZH1-p65 axis to mediate angiogenesis and alveolarization of BPD mice.

**Conclusion:** Altogether, the present study revealed that HDAC3 could regulate the EZH1-p65-Pgf axis through miR-17 in the miR-17-92 cluster in the pulmonary angiogenesis and alveolarization of BPD mice.

## Background

Bronchopulmonary dysplasia (BPD) is a common chronic lung disease characterized by arrested alveolar development or loss of alveoli which commonly occurred in premature infants. BPD was associated with increased risks of morbidity and mortality, as well as neurodevelopmental outcomes among prematurely born infants [1, 2]. Risk factors responsible for BPD include pre- and postnatal infections, hyperoxia, as well as mechanical ventilation, which not only influence the interacted function of pro- and anti-inflammatory proteins, but also the extra alterations of signaling pathways which is closely related to the imbalance of growth factor [3]. Although extensive approaches have alleviated the survival of preterm infants in neonatal care, the effectiveness of the currently available therapeutic strategies in reducing the incidence and severity of BPD remains very limited [4]. Therefore, it is important to determine the specific molecular mechanism of BPD in order to identify potential novel biomarkers relevant to BPD.

Histone deacetylases (HDACs) are chromatin-modifying enzymes that have emerged as the important targets for treating various diseases due to their powerful regulatory role in physiological and pathological settings [5]. HDAC3, a member of the HDAC superfamily, has been reported to be highly expressed in adenocarcinoma of the lung and closely associated with the poor prognosis of this disease [6]. Additionally, HDAC3 has been shown to play a pivotal role in cellular proliferation, apoptosis as well as transcriptional repression, and may serve as a negative regulator of angiogenesis [7]. However, the regulatory function of HDAC3 in BPD remains unknown. Evidence has shown that the knockdown of HDAC3 could upregulate the expression of microRNA (miR)-15a/16-1, thereby suppressing lung cancer cell growth and colony formation [8]. miRNAs are a class of small noncoding RNAs that regulate gene expression at the post-transcriptional level, leading to mRNA degradation or inhibition of protein translation [9]. The miR-17-92 cluster (miR-17, miR-18a, miR-19a, miR-19b, miR-20a, and miR-92) has been reported to be poorly expressed in lung tissues of BPD infants, while the downregulation of miR-17-92 cluster was closely related to the development of BPD [10]. It has been shown that miR-17 could regulate the expression of EZH1 which involved in the mediation of drug resistance in non-small cell lung cancer, Besides, the binding of EZH1 to the p65 transcription factor could promote the transcription of downstream target genes [11]. It has also been reported that p65 could directly bind to the promoter region of placental growth factor (Pgf)/Plgf and transcriptionally activates the expression of Pgf/Plgf [12]. Pgf has been found to be involved in the regulation of BPD, as well as acute and chronic lung injury in newborn infants [13]. Therefore, it was hypothesized that EZH1 could promote p65-activated transcription of Pgf in BPD mice. In the present work, we planned to explore the potential effects of HDAC3 regulating miR-17 in the miR-17-92 cluster on the angiogenesis and alveolarization of BPD *via* EZH1-p65-Pgf axis, which could serve as a potential target to treat BPD.

## Materials and methods

### Ethics statement

All animal experiments were conducted in strict accordance with the Guide to the Management and Use of Laboratory Animals issued by the National Institutes of Health. The protocol of animal experiments was approved by the Institutional Animal Care and Use Committee of Affiliated Hangzhou First People’s Hospital, Zhejiang University School of Medicine.

### Hyperoxia-induced BPD mouse model establishment

Thirty C57Bl/6J newborn mice with an average weight of 1.36 g ± 0.09g were randomly divided into one litter (equal numbers of mice per litter). The mice following BPD modeling were exposed to 85% oxygen for postnatal day 1 (P1) - P14, while the control mice with normal lung development were exposed to 21% oxygen. Male and female animals were used because no gender bias was found in the study on perturbations to lung development of C57Bl/6J mice in response to hyperoxia. The care dam was rotated every 24 h under normal hypoxia and hyperoxia in order to limit oxygen toxicity. Mice in P3 (before the formation of large alveoli), P5 (peak of large alveolar formation), or P14 (the stage of massive alveolarization) were euthanized by intraperitoneal injection of excessive pentobarbital (500 mg/kg), and the lungs of mice were then extracted for further analyses. The mice were then randomly assigned into the control group (normal air-treated C57B1/6J newborn mice), the BPD group (hyperoxia-induced BPD mice), the BPD + HDAC3^−/−^ group (BPD mice treated with HDAC3 knockout) [14], the hyperoxia + HDAC3^−/−^ + antagomir NC group (hyperoxia-induced BPD mice with HDAC3 knockout and injected with antagomir NC), and the hyperoxia + HDAC3^−/−^ + miR-17-antagomir group (hyperoxia-induced BPD mice with HDAC3 knockout and injected with miR-17-antagomir), with ten mice in each group.

### Cell culture and grouping

Primary lung fibroblasts were isolated from C57B1/6J mice. Briefly, the lungs were injected with approximately 500 μL of preheated collagenase type I (2 mg/mL) at 37°C and then excised from adult female C57BL/6J mice after euthanized by isoflurane inhalation. The lung tissues were then placed in a 50 mL tube containing 25 mL of preheated collagenase type I and treated at 70 rpm for 1 h at 37°C using an orbital rotator (Unimax 1010) with gentle agitation. The lungs were then minced using sterile scissors and the tissue suspension was dispersed by using a 20G syringe needle. The cell suspension was then injected through a 40 mL filter into a new 50 mL tube, followed by centrifugation at 120 g at 4°C for 8 min, with the supernatant discarded. The cell pellet was then resuspended in preheated (37°C) high-glucose Dulbecco’s modified Eagle’s medium (DMEM) containing 10% (v/v) fetal bovine serum (FBS), 100 U/mL penicillin (ThermoFisher Scientific, Waltham, MA, USA) and 100 μg/mL streptomycin (ThermoFisher Scientific), followed by inoculation into a T-75 cell culture flask (1 flask per lung), then by passage in low-glucose DMEM containing 10% (v/v) FBS, 100 U/mL penicillin, and 100 μg/mL streptomycin. The cells used in this study were primary isolated and cultured lung fibroblasts [15].

The lung fibroblasts were cultured in a 6-well plate at a cell concentration of 2 × 10^5^ cells/per tube. When the cells reached 80% confluence, transfection was performed according to the manufacturer’s instructions of the lipofectamine 2000 specifications (11668-019, Invitrogen, New York, California, USA). The fibroblasts were transfected with following plasmids: short hairpin (sh) RNA-negative control (NC), sh-HDAC3, inhibitor-NC, miR-17-inhibitor, sh-HDAC3 + inhibitor-NC, sh-HDAC3 + miR-17-inhibitor, overexpressed (oe)-NC, oe-EZH1, sh-NC, sh-p65 and oe-EZH1 + sh-p65 (Guangzhou RiboBio Co., Ltd., Guangzhou, Guangdong, China). After 12 h of transfection, the cells were cultured for 48 h at 37°C under 5% CO_2_ for subsequent RNA extraction and other relevant experiments.

### Hematoxylin-eosin (HE) staining

The left lung tissues of mice in different groups were fixed in 10% neutral formaldehyde solution for more than 24 h, paraffin-embedded, dewaxed twice with xylene (10 min/time), then rehydrated with 100% ethanol for 5 min, 90% ethanol for 2 min, 70% ethanol for 2 min, washed with distilled water for 2 min and stained with hematoxylin for 7 min. After washing for 10 min with tap water to remove the excess dye solution, the samples were then washed with distilled water again and subsequently treated with 95% ethanol for 5 s, stained with eosin for 1 min, hydrated using gradient ethanol (100%, 95%, 75%, 50%, twice, 2 min/time), cleared twice by xylene (5 min/time), and air-dried. The lung tissues were then blocked with neutral resin in a fume hood and the morphological changes of lung tissues were observed under an optical microscope.

### Immunohistochemistry

Left lung tissues were taken from the euthanized mice, fixed in 4% paraformaldehyde, embedded in paraffin, serially sectioned at a thickness of 4 μm and routinely dewaxed. The streptavidin-peroxidase (SP) method was routinely conducted. Briefly, antigen retrieval was performed by microwave heating. The sections were blocked by the addition of normal goat serum blocking solution. BPD group was used as a NC. The staining was conducted with the HistostainTM SP-9000 immunohistochemical staining kit (Zymed Laboratories Inc., South San Francisco, CA, USA). The sections were then probed at 4°C overnight with the following primary antibodies purchased from Abcam Inc. (Cambridge, UK): rabbit anti-EZH1 (ab64850, 1: 100), rabbit anti-NF-κB p65 (ab16502, 1: 100), and rabbit anti-Pgf (ab230516, 1: 100). After rinsing with PBS, the sections were immediately probed with mouse anti-rabbit secondary antibody (ab6728, 1: 1000, Abcam) at 37°C for 30 min. After PBS washing, the horseradish peroxidase (HRP)-labeled working solution was added to the sections and incubated, followed by diaminobenzidine (DAB) development for 5 to 10 min. The staining time was adjusted under a microscope and the sections were sealed using gum after 1 min of hematoxylin counterstaining. Then, five representative high-power fields were selected for cell observation and counting. The cytoplasm stained brown or yellow were indicative of positive cells.

### Immunofluorescence staining of microvessel density (MVD)

The lung tissue sections were deparaffinized, rehydrated, and stained with rabbit polyclonal antibody to Von Willebrand Factor (vWF, ab11713, 1: 100, Abcam). The number of blood vessels (20-50 μm diameter) of each high power field (HPF) was counted in five randomly selected non-overlapping parenchymal regions of the lung tissue sections of the animals (n = 6). The detailed procedures were carried out as previously described [16].

### Reverse transcription-quantitative polymerase chain reaction (RT-qPCR)

Lung fibroblasts in each group were collected and lysed using the TRIzol kit (Invitrogen). Then, the total RNA was extracted from cells and tissue samples. The quality and concentration of RNA were measured using an ultraviolet-visible spectrophotometer (ND-1000, Nanodrop, Wilmington, DE, USA). An amount of 400 ng of the extracted RNA was subjected to reverse transcription by using a PrimeScript RT Reagent Kit (Takara, Shiga, Japan). With complementary DNA (cDNA) as a template, a fluorescent quantitative PCR was carried out in accordance with the manufacturer’s instructions of a SYBR® Premix Ex Taq^TM^ II (Tli RNaseH Plus) kit (Takara). The primers were synthesized by RiboBio Company and were shown in Table 1. Glyceraldehyde-3-phosphate dehydrogenase (GAPDH) or U6 was used as a loading control gene. The fold changes were calculated using the relative quantification (the 2^−ΔΔCt^ method): ΔΔCT = Ct_(target gene)_ -Ct _(internal reference gene)_.

**Table 1.**
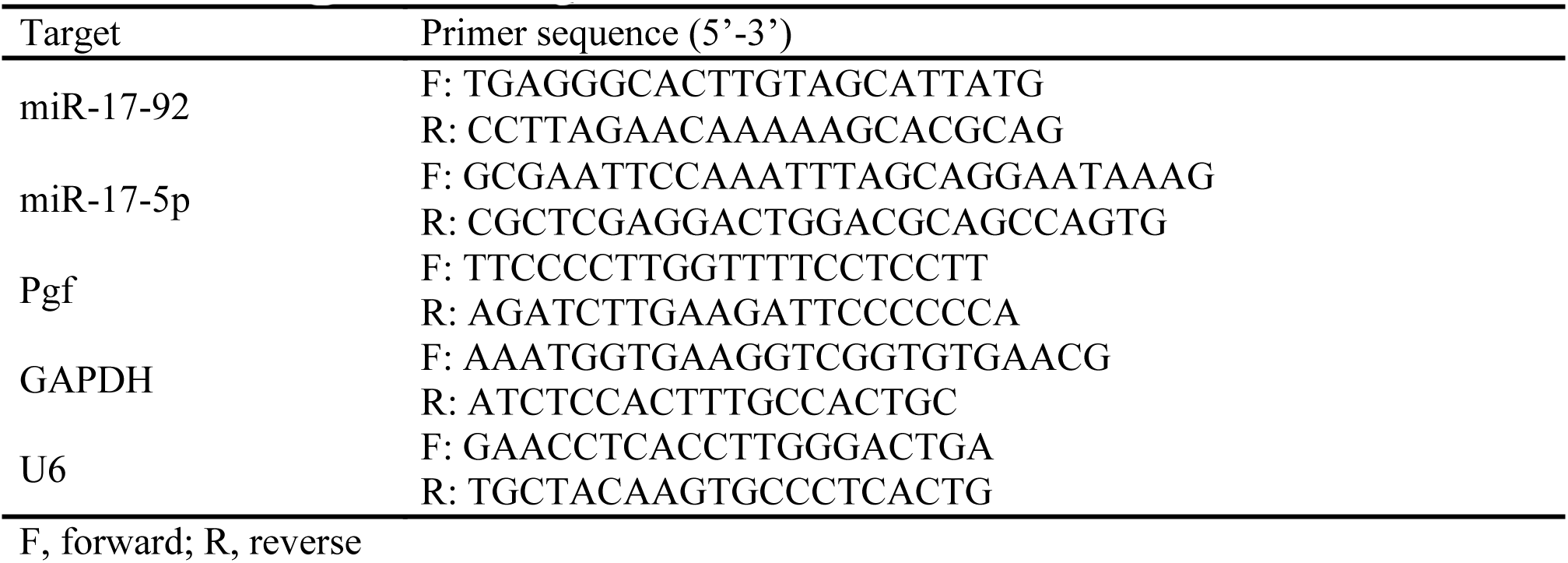
Primer sequences for RT-qPCR

### Western blot analysis

Lung fibroblasts were washed with PBS, lysed with cell lysis buffer (C0481, Sigma-Aldrich, St. Louis, MO, USA) and incubated at 4°C for 30 min. Next, the cell lysate was collected into 1.5 mL Eppendorf tubes and centrifuged at 12,000 g at 4°C for 5 min, after which the supernatant was collected. The concentration of protein was measured using the bicinchoninic acid protein assay kit (Shanghai Beyotime Biotechnology Co. Ltd., Shanghai, China). Subsequently, 20 μg of protein was separated by 10% sodium dodecyl sulfate-polyacrylamide gel electrophoresis (SDS-PAGE) (Millipore, Billerica, MA, USA) and transferred onto a polyvinylidene fluoride membrane (Millipore). The membrane was then blocked with 5% skimmed milk powder for 1 h and probed with the following Tris-buffered saline Tween-20 (TBST) diluted primary antibodies purchased from Abcam: rabbit polyclonal antibody to HDAC3 (ab7030, 1: 1000), EZH1 (ab64850, 1: 250), p65 (ab19870, 1: 500) and Pgf (ab230516, 1: 1000) at 4°C overnight, followed by washing 3 times with TBST and then probed with HRP-labeled secondary antibody goat anti-mouse or goat anti-rabbit (HS101, 1: 1000; TransGen Biotech Co., Ltd., Beijing, China) at room temperature for 1 h. Following 6 rinses with TBST, the immunocomplexes on the membrane were visualized using an enhanced chemiluminescence kit (Shanghai Baoman Biotechnology Co., Ltd., Shanghai, China). With GAPDH (ab37168, 1: 1000, Abcam) as a loading control, the band intensities were quantified using Image J analysis software.

### Chromatin immunoprecipitation (ChIP) assay

ChIP assay was used to detect the enrichment of HDAC3 in the promoter region of the miR-17-92 cluster and that of p65 in the Pgf promoter region using the EZ-Magna ChIP TMA kit (Millipore). The lung fibroblasts of each group in logarithmic growth phase were cross-linked and cultured with 1% formaldehyde for 10 min, then terminated by adding 125 mM glycine at room temperature for 5 min. The cells were then washed twice with pre-chilled PBS and centrifuged at 2000 rpm for 5 min after which the supernatant was collected. Next, the cells were resuspended in cell lysate [150 mM Sodium chloride (NaCl), 50 mM Tris (pH 7.5), 5 mM ethylenediamine tetraacetic acid (EDTA), 0.005% NP40, 0.01% Triton X-100)] to prepare a final concentration of 2 × 10^6^ cells/200 mL. The cells were then added with protease inhibitor mixture, centrifuged at 5000 rpm for 5 min, resuspended in nuclear separation buffer, lysed in ice water bath for 10 min and sonicated to obtain a 200-1000 bp chromatin fragment. The supernatant was then aspirated following centrifugation at 14,000 g for 10 min at 4°C. Then, 100 μL of the supernatant (DNA fragment) in each group was collected and added with 900 μL of ChIP Dilution Buffer, 20 μL of 50 × pseudoisocyanine (PIC) and 60 μL of ProteinA agarose/salmon sperm DNA, followed by mixing at 4°C for 1 h, allowed to stand at 4°C for 10 min and centrifuged at 700 rpm for 1 min. The supernatant was then collected, 20 μL of which was taken as the Input. In the experimental groups, the supernatant was separately incubated with 1 μL of rabbit anti-HDAC3 (ab7030, 1: 500, Abcam) and rabbit anti-p65 (ab19870, 1: 100, Abcam). The supernatant in the NC groups was incubated with 1 μL of rabbit anti-immunoglobulin G (IgG) (ab172730, Abcam) and each tube was added with 60 μL of ProteinA agarose/salmon sperm DNA and incubated at 4°C for 2 h. Following standing for 10 min, the samples were centrifuged at 700 rpm for 1 min and the supernatant was discarded. The precipitate was washed with 1 mL of low salt buffer, high salt buffer, LiCl solution, and Tris-EDTA (TE) buffer (twice) respectively. Each tube was eluted twice with 250 mL of ChIP Wash Buffer and decrosslinked with 20 mL of 5 M NaCl, after which the DNA was recovered and the promoter sequences of miR-17-92 and Pgf in the complex were quantified by RT-qPCR.

### Dual-luciferase reporter gene assay

The constructed 3’ untranslated region (3’UTR) luciferase vector of EZH1 was cloned into the pMIR reporter plasmid by PCR amplification of the potential binding site. For dual-luciferase reporter gene assay, based on Lipofectamine 2000 (Invitrogen), lung fibroblasts containing the EZH1 3’UTR and miR-17-5p mimics and inhibitors were transiently transfected into the pMIR reporter vector. After 48 h of transfection, reporter gene activity was measured using a dual-luciferase assay system (Promega, Madison, WI, USA). Renilla luciferase activity was used to normalize transfection efficiency. The promoter activity of HDAC for miR-17-92 cluster was described previously [14]. The effect of p65 on the promoter activity of Pgf was described previously [17].

### Statistical analysis

Statistical analyses were conducted using SPSS 21.0 software (IBM Corp. Armonk, NY, USA). Measurement data were expressed as mean ± standard deviation. Confirming to normal distribution and homogeneity of variance, the data between two groups with the unpaired design were compared using unpaired *t*-test. Data among multiple groups were analyzed by one-way analysis of variance (ANOVA), followed by a Tukey’s post-hoc test. The difference in *p*-value of *p* < 0.05 was considered as statistically significant.

## Results

### HDAC3 was involved in the regulation of BPD

Previous literature has reported that HDAC3 was participated in the remodeling and expansion of distant alveolar vesicles into primitive pulmonary alveolus by regulating the miR-17-92 cluster [14], which commonly constitutes to the occurrence of BPD in premature infants [15]. Based on the aforementioned evidence, it was suggested that HDAC3 may be involved in the progression of BPD. Therefore, the molecular mechanism of HDAC3 involved in BPD was explored by establishing a hyperoxia-induced mouse model of BPD. The degree of lung injury in mice was firstly evaluated by HE staining. The results showed that the degree of lung injury in the BPD mice was notably higher than that in the control mice (Fig. 1A). The degree of alveolarization in BPD mice was subsequently examined and the results showed that the mean linear intercept (MLI) of alveoli in BPD mice was significantly higher than that of the control mice, while the number of alveoli was less than that of the control mice (Fig. 1B). Furthermore, the MVD in lung tissues was examined by immunofluorescence assay, and the results suggested that the MVD of BPD mice was significantly lower than that of the control mice (Fig. 1C). The expression of HDAC3 in lung tissues of BPD mice was assessed by Western blot analysis, which demonstrated that the expression of HDAC3 in lung tissues of BPD mice was markedly higher than that of control mice (Fig. 1D). Next, the HDAC3 gene was knocked out in BPD mice, and it was verified that the degree of lung injury in BPD + HDAC3^−/−^ mice was significantly reduced (Fig. 1A), the degree of alveolarization was notably lower (Fig. 1B), as well as the MVD was higher when compared to BPD mice (Fig. 1C). The expression of HDAC3 in lung tissues of BPD + HDAC3-/- mice was also significantly lower than that in the BPD mice as detected by Western blot analysis (Fig. 1D). These results demonstrated that HDAC3 was involved in the regulation of alveolarization and angiogenesis in BPD.

**Fig. 1.**
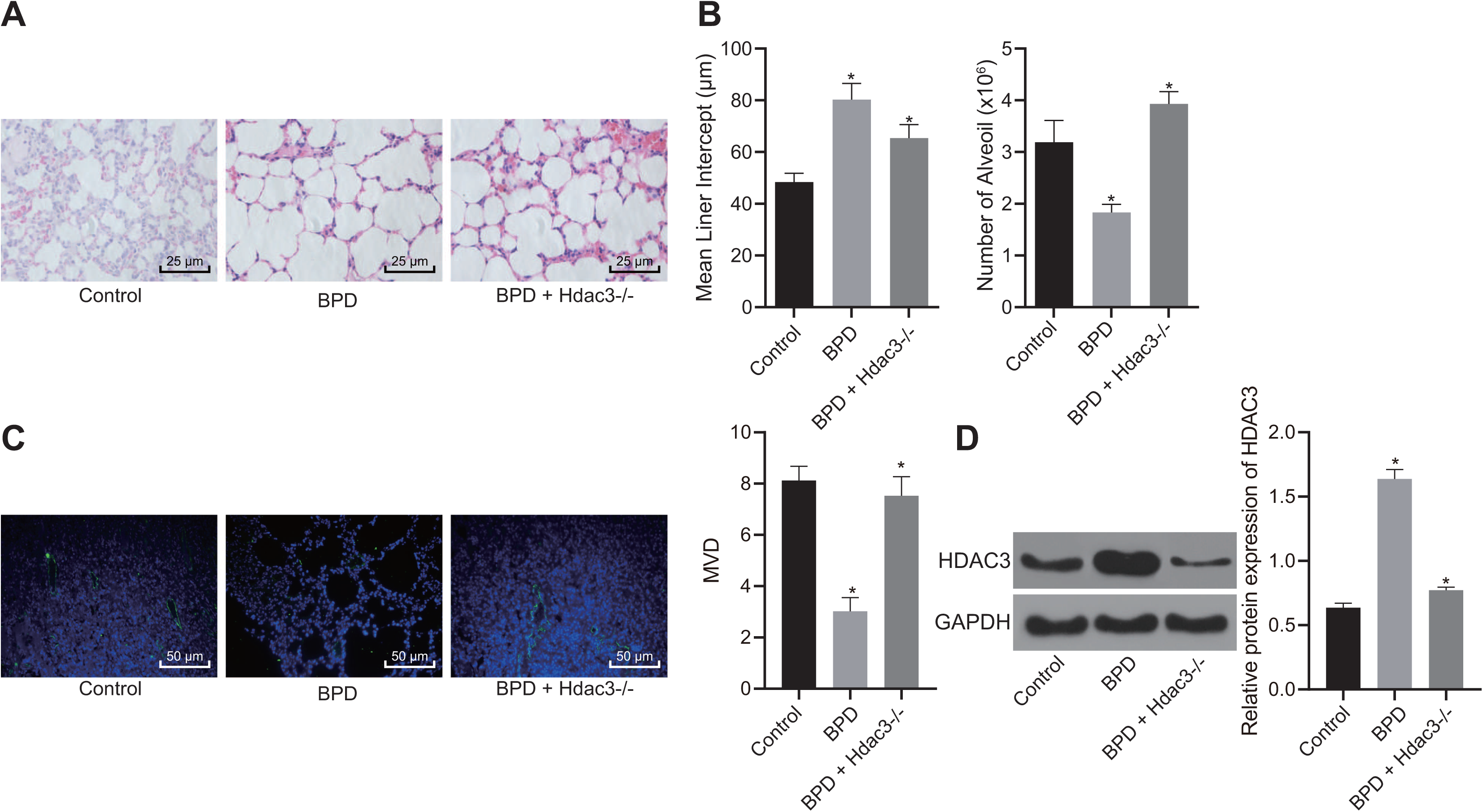
HDAC3 is implicated in the regulation of alveolarization and angiogenesis in BPD. A, The degree of lung injury of mice in each group detected by HE staining (× 400). B, The degree of alveolarization and MLI of alveoli in BPD mice. C, MVD in lung tissues of mice in each group assessed by immunofluorescence assay (× 200). D, HDAC3 protein expression in lung tissues of mice in each group detected by Western blot analysis. * *p* < 0.05. The above data were measurement data and expressed as mean ± standard deviation. Data among multiple groups were analyzed by one-way ANOVA, followed by Tukey’s post-hoc test. n = 10 for each mouse group.

### HDAC3 inhibits the expression of miR-17-92 cluster in BPD

RT-qPCR was used to detect the expression of miR-17-92 cluster (miR-17, miR-18a, miR-19a, miR-19b, miR-20a, miR-92) in control mice, BPD mice and BPD + HDAC3^−/−^ mice. The results showed that the expression of the miR-17-92 cluster in BPD mice was significantly lower than that in control mice, but demonstrated increased expression of miR-17-92 cluster after HDAC3 knockout (Fig. 2A). Furthermore, the primary lung fibroblasts were isolated from BPD mice and stably transfected with sh-NC and sh-HDAC3. The protein level of HDAC3 was detected by western blot analysis. Results demonstrated that the protein level of HDAC3 was notably decreased following silencing of HDAC3 expression (Fig. 2B). The expression of miR-17-92 cluster was then detected in the cells, and the results indicated that the expression of miR-17-92 cluster in sh-HDAC3-treated cells was markedly higher than those of sh-NC-treated cells (Fig. 2C), implying that HDAC3 may regulate miR-17-92 cluster in BPD. To further elucidate the enrichment of HDAC3 in the promoter region of miR-17-92 cluster, ChIP assay showed that HDAC3 was enriched in the miR-17-92 cluster promoter, while silencing of HDAC3 resulted in the opposite trend (Fig. 2D). Furthermore, the effect of HDAC3 on the luciferase activity of miR-17-92 promoter was examined by dual-luciferase reporter gene assay. The results showed that silencing of HDAC3 expression resulted in inhibited luciferase activity of miR-17-92 promoter (Fig. 2E). Based on the aforementioned results, it was suggested that HDAC3 could regulate the expression of miR-17-92 cluster in BPD.

**Fig. 2.**
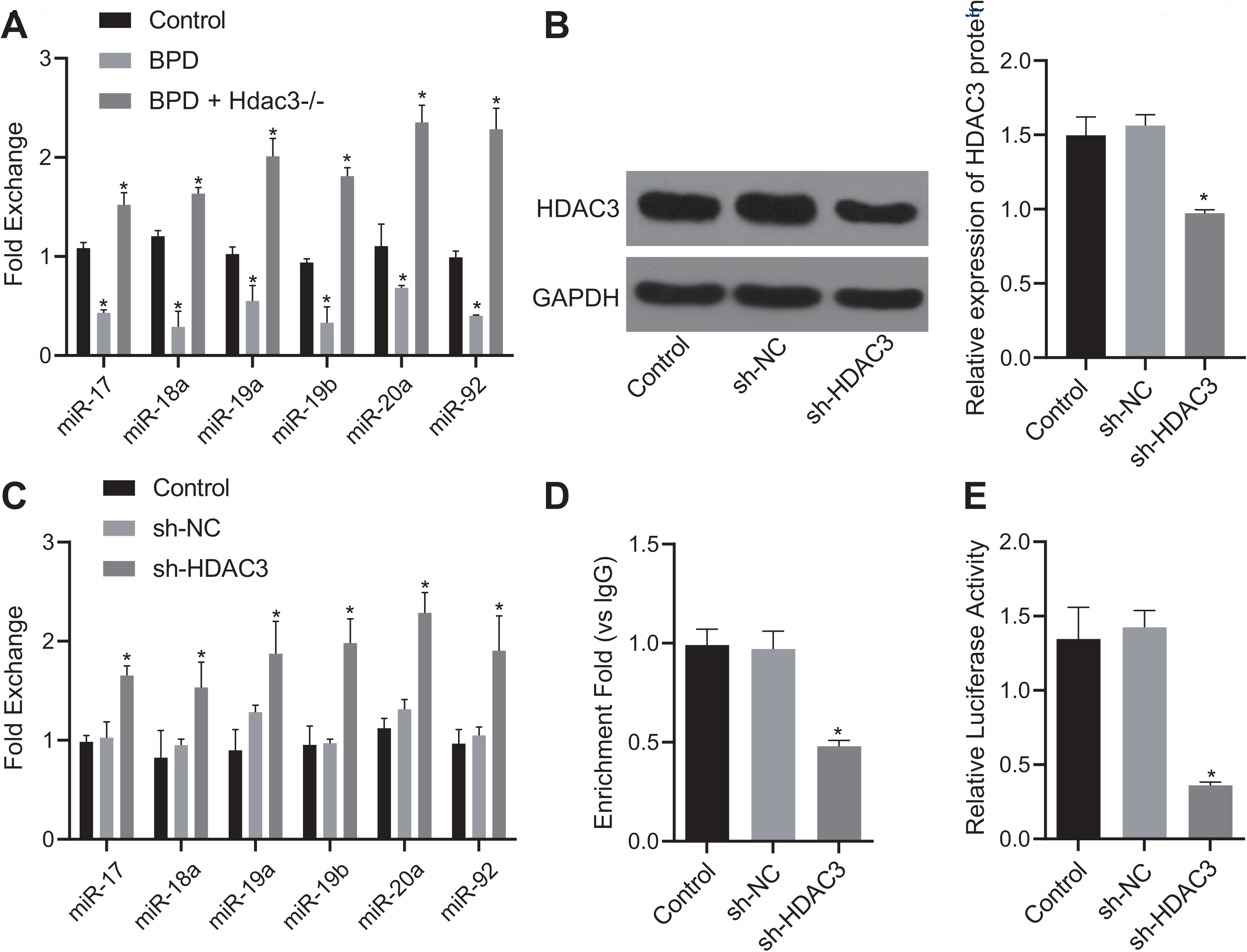
HDAC3 suppresses the expression of the miR-17-92 cluster in BPD. A, The expression of miR-17-92 cluster in lung tissues of different groups of mice detected by RT-qPCR. B, Western blot analysis of HDAC3 protein expression after silencing of HDAC3 expression. C, The expression of miR-17-92 cluster in cells following HDAC3 silencing detected by RT-qPCR. D, The enrichment of HDAC3 in the promoter region of miR-17-92 cluster in cells following different transfection. E: Luciferase activity of miR-17-92 cluster promoter detected by dual-luciferase reporter gene assay. * *p* < 0.05. The above data were measurement data and expressed as mean ± standard deviation. Data among multiple groups were analyzed by one-way ANOVA, followed by Tukey’s post-hoc test. n = 10 for each mouse group.

### EZH1 was a target gene of miR-17 in miR-17-92 cluster

To further explore the underlying mechanism of miR-17 in the miR-17-92 cluster in the regulation of BPD, the Targetscan website was employed to predict whether miR-17 could bind to EZH1. It was verified that miR-17 could bind to EZH1 (Fig. 3A). Previous study has been reported that miR-17 could be involved in the regulation of drug resistance in non-small cell lung cancer *via* targeting and regulating EZH1 expression [11]. To further explore the regulatory mechanism of miR-17 and EZH1 in alveolarization, lung fibroblasts were stably transfected with inhibited and overexpressed miR-17 in BPD. RT-qPCR was performed to detect miR-17 expression in cells, and the results showed that inhibited miR-17 resulted in decreased miR-17 expression, while overexpression of miR-17 resulted in increased miR-17 expression (Fig. 3B). The binding of miR-17 to EZH1 was then examined by a dual-luciferase reporter gene assay, and the results revealed that overexpression of miR-17 resulted in inhibited luciferase activity of EZH1 3’UTR, while suppression of miR-17 resulted in promoted luciferase activity of EZH1 3’UTR (Fig. 3C). Results from RT-qPCR and Western blot analysis demonstrated that overexpressed miR-17 resulted in inhibited EZH1 expression, whereas inhibited miR-17 resulted in promoted EZH1 expression (Fig. 3D). To further investigate whether HDAC3 could affect EZH1 expression through miR-17 in lung fibroblasts in BPD, the expression of EZH1 in fibroblasts transfected with sh-HDAC3, or with both HDAC3 silencing and miR-17 inhibitor was examined. The results verified that the expression of EZH1 was inhibited by silencing of HDAC3 expression, but it could be restored by the co-transfection of HDAC3 silencing and miR-17 inhibitor (Fig. 3E). The above results showed that miR-17 was involved in the regulation of EZH1 by modulating HDAC3 in BPD.

**Fig. 3.**
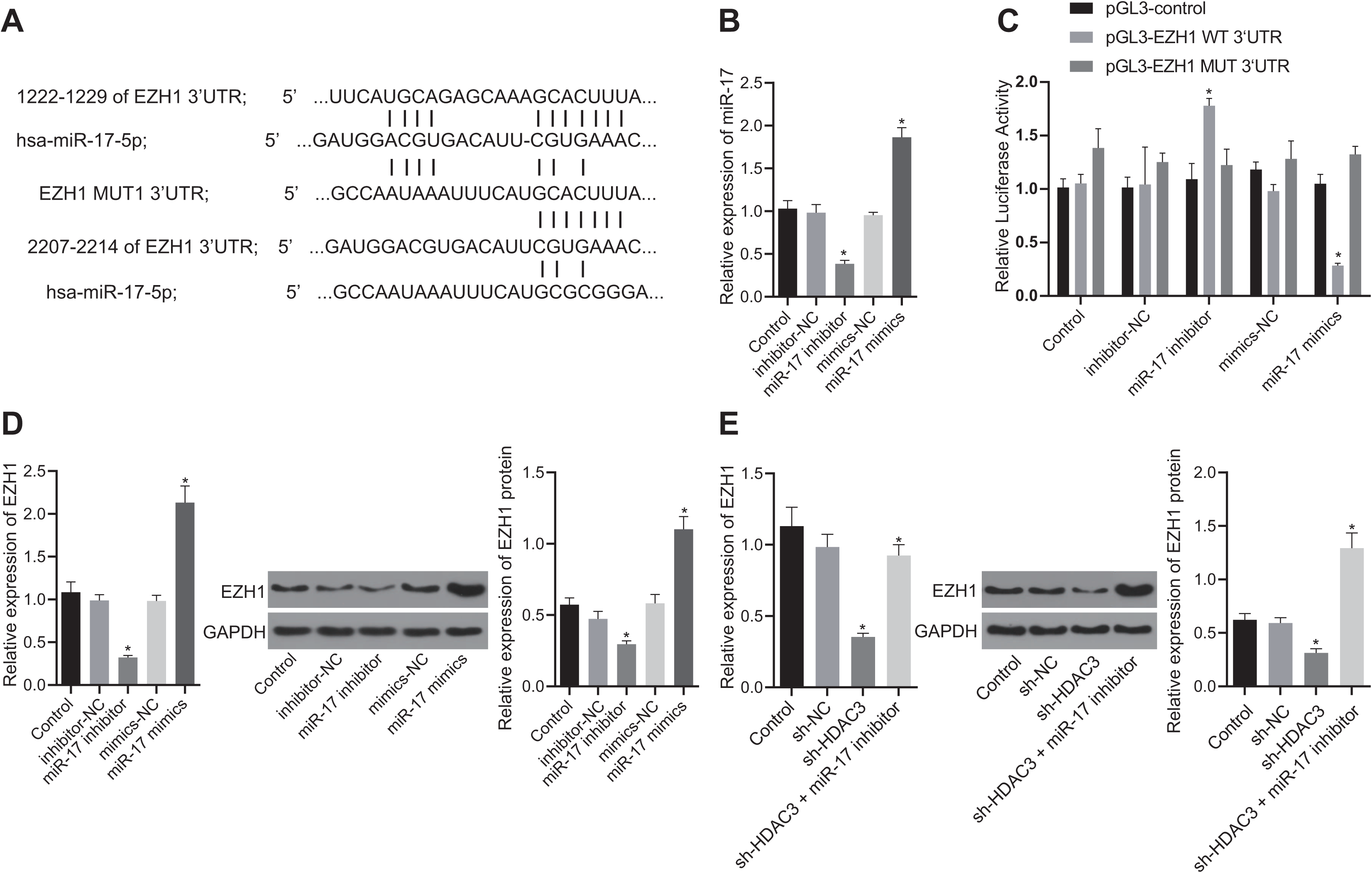
miR-17 in the miR-17-92 cluster targets EZH1 and regulates its expression in primary lung fibroblasts of BPD mice. A, The binding site of miR-17 and EZH1 predicted by Targetscan website. B, The expression of miR-17 in cells transfected with miR-17 mimic and miR-17 inhibitor detected by RT-qPCR. C, The binding of miR-17 to EZH1 detected by dual-luciferase reporter gene assay. D, The expression of EZH1 in cells transfected with miR-17 mimic and miR-17 inhibitor detected by RT-qPCR and Western blot analysis. E, The expression of EZH1 in fibroblasts transfected with HDAC3 silencing, or both HDAC3 silencing and miR-17 inhibitor detected by RT-qPCR and Western blot analysis. * *p* < 0.05. The above data were measurement data and expressed as mean ± standard deviation. Data among multiple groups were analyzed by one-way ANOVA, followed by Tukey’s post-hoc test. n = 10 for each mouse group.

### EZH1 activates p65 transcription factor to enhance Pgf expression

According to previous literature, EZH1 could bind to p65 transcription factor to promote the transcription of downstream target genes [11], while in human Plgf promoter region, p65 has been reported to bind directly to the promoter region of Pgf/Plgf and transcriptionally activates Pgf/Plgf [12]. Pgf has been demonstrated to be involved in the regulation of BPD [13]. Therefore, it was hypothesized that EZH1 may promote p65 to activate the transcription of Pgf in BPD mice. Initially, the binding sites of p65 and Pgf were obtained (Fig. 4A). The expression of EZH1 in fibroblasts transfected with oe-EZH1 was detected by RT-qPCR and Western blot analysis. It was shown that the expression of EZH1 was significantly up-regulated after the transfection with oe-EZH1 in fibroblasts of BPD mice (Fig. 4B). The binding of EZH1 to p65 was further examined by immunoprecipitation (IP) assay, and the results showed that the overexpression of EZH1 promoted the binding of EZH1 to p65 (Fig. 4C). Besides, results from the ChIP assay revealed that the overexpression of EZH1 promoted the enrichment of p65 in the Pgf promoter region (Fig. 4D). The non-fibroblasts of BPD mice that stably transfected with silencing of p65 were constructed and the expression of p65 was detected. The results showed that silencing of p65 expression resulted in inhibited expression of p65 in non-fibroblasts of BPD mice (Fig. 4E). The expression of Pgf was then detected in silencing p65-treated cells, and the results suggested that silencing of p65 expression resulted in inhibited expression of Pgf (Fig. 4F). Moreover, it has been indicated that EZH1 may be involved in the regulation of Pgf expression. Next, Pgf expression was detected in oe-EZH1-treated cells. The overexpression of EZH1 resulted in promoted Pgf expression, and the expression of Pgf was rescued in cells after co-transfection of sh-p65 and oe-EZH1 (Fig. 4G). The above-mentioned data indicated that EZH1 could participate in the regulation of BPD by recruiting p65 to promote Pgf expression.

**Fig. 4.**
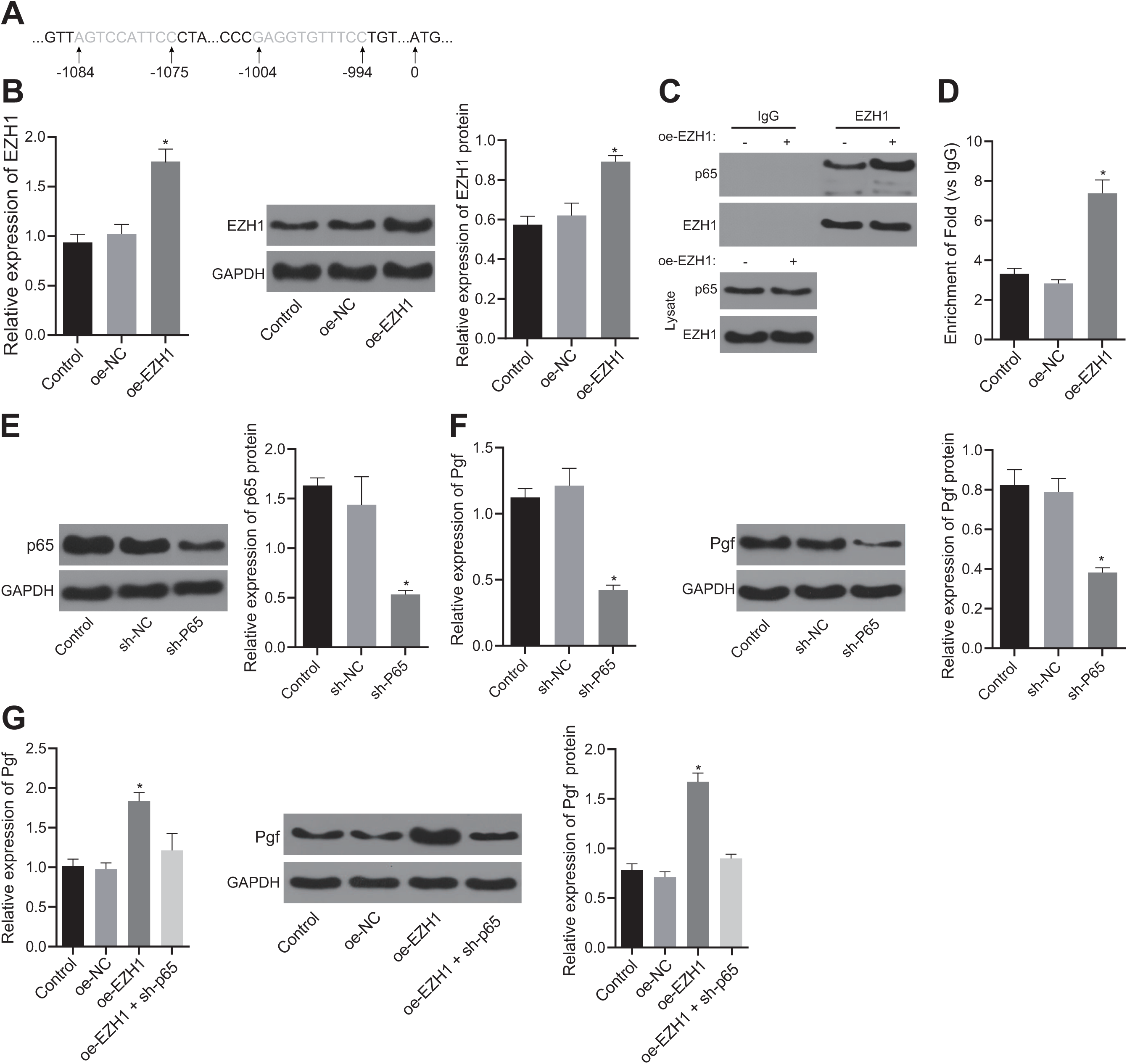
EZH1 promotes the expression of Pgf *via* p65 transcription factor stimulation. A, The binding sites between p65 and Pgf. B, The expression of EZH1 in cells transfected with oe-EZH1 detected by RT-qPCR and Western blot analysis. C, The binding of EZH1 and p65 detected by IP assay. D, The enrichment of p65 in the Pgf promoter region in oe-EZH1-treated cells detected by ChIP assay. E, The expression of p65 in p65 silencing-treated cells detected by Western blot analysis. F, The expression of Pgf in p65 silencing-treated cells detected by RT-qPCR and Western blot analysis. G, The expression of Pgf in fibroblasts transfected with oe-EZH1, or both oe-EZH1and sh-p65 detected by RT-qPCR and Western blot analysis. * *p* < 0.05. The above data were measurement data and expressed as mean ± standard deviation. The data between the two groups with unpaired design were compared by unpaired *t*-test. Data among multiple groups were analyzed by one-way ANOVA, followed by Tukey’s post-hoc test. n = 10 for each mouse group.

### HDAC3 regulates Pgf expression *via* miR-17-EZH1-p65 axis

To further explore the regulatory role of HDAC3 on Pgf expression in BPD, the expression of Pgf was initially assessed by RT-qPCR and western blot analysis in cells transfected with silencing of HDAC3 expression and inhibited miR-17. The corresponding results suggested that the silencing of miR-17 resulted in promoted Pgf expression, whereas silencing of HDAC3 resulted in inhibited Pgf expression, while Pgf expression was rescued in cells after co-transfection of sh-HDAC3 and miR-17 inhibitor (Fig. 5A). According to our previous results, HDAC3 regulated the expression of EZH1 through miR-17, therefore it was hypothesized that HDAC3 could promote the transcription of Pgf *via* miR-17-EZH1-p65 axis. ChIP assay was used to detect the enrichment of p65 in the Pgf promoter region in cells transfected with sh-HDAC3, miR-17 inhibitor, or both. The results showed that silencing of HDAC3 inhibited the enrichment of p65 in the Pgf promoter region while silencing of miR-17 promoted the enrichment of p65 in the Pgf promoter region. Besides, the co-transfection of sh-HDAC3 and miR-17 inhibitor relatively restored the enrichment of p65 in the Pgf promoter region (Fig. 5B). These results demonstrated that HDAC3 could enhance the transcription and expression of Pgf *via* miR-17-regulated EZH1 and subsequently promote the recruitment of p65 in the Pgf promoter region.

**Fig. 5.**
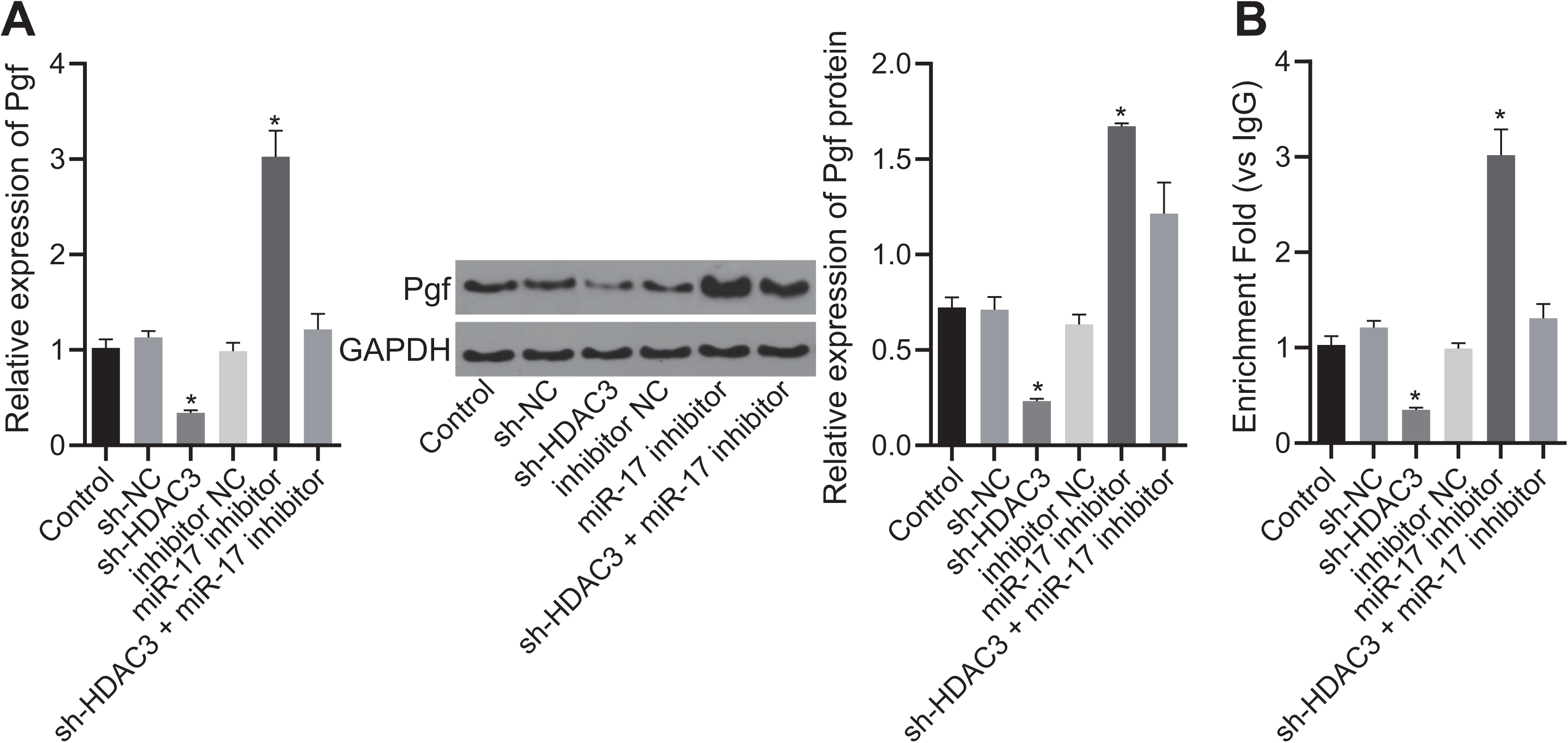
HDAC3 potentiates the transcription and the expression of Pgf *via* the miR-17-EZH1-p65 axis. A, The expression of Pgf detected by RT-qPCR and Western blot analysis. B, The enrichment of p65 in Pgf promoter region detected by ChIP assay. * *p* < 0.05. The above data were measurement data and expressed as mean ± standard deviation. Data among multiple groups were analyzed by one-way ANOVA, followed by Tukey’s post-hoc test. n = 10 for each mouse group.

### HDAC3 regulates Pgf *via* miR-17 in miR-17-92 cluster and promotes BPD development

To further demonstrate the regulatory role of HDAC3 on Pgf in BPD mice *via* the miR-17-EZH1-p65 axis *in vivo*, hyperoxia + HDAC3^−/−^ + miR-17-antagomir mice were constructed through injection of miR-17 antagomir into hyperoxia + HDAC3^−/−^ mice. The expression of miR-17 in mice following different treatment was initially detected by RT-qPCR. The results showed that miR-17 expression in hyperoxia + HDAC3^−/−^ mice was significantly higher than that in hyperoxia-induced BPD mice, but relatively decreased in hyperoxia + HDAC3^−/−^ + miR-17-antagomir-treated mice, which was similar to that in the BPD mice (Fig. 6A). Furthermore, the degree of lung injury in mice was detected by HE staining. It was demonstrated that the degree of lung injury in hyperoxia + HDAC3^−/−^-treated mice was notably improved when compared to the hyperoxia-induced BPD mice. The degree of lung injury in hyperoxia + HDAC3^−/−^ + miR-17-antagomir-treated mice was similar to that in the hyperoxia-induced BPD mice (Fig. 6B). The alveolar number and MLI of the mice were subsequently detected to evaluate the degree of alveolarization in mice. Results demonstrated that the MLI of hyperoxia + HDAC3^−/−^ mice-treated was significantly reduced, and that of the MLI of hyperoxia + HDAC3^−/−^ + miR-17-antagomir mice was similar to that of hyperoxia-induced BPD mice (Fig. 6C). The number of alveolar in hyperoxia + HDAC3^−/−^-treated mice was notably higher than that in the BPD mice, and the number of alveoli in hyperoxia + HDAC3^−/−^+ miR-17-antagomir-treated mice was similar to that in the hyperoxia-induced BPD mice (Fig. 6D). The MVD of lung tissue in each group was detected by immunofluorescence assay. It was shown that the MVD of hyperoxia + HDAC3^−/−^-treated mice was markedly higher than that of BPD mice, and the MVD of hyperoxia + HDAC3^−/−^ + miR-17-antagomir-treated mice was similar to that of the hyperoxia-induced BPD mice (Fig. 6E). Besides, it was shown that HDAC3 could regulate the alveolarization and angiogenesis *via* miR-17 in BPD mice. Furthermore, the expression of EZH1, p65, and Pgf in lung tissue of mice in different groups was examined by immunohistochemistry. The results showed that the expression of EZH1, p65, and Pgf in hyperoxia + HDAC3^−/−^-treated mice was significantly lower than that in BPD mice, whereas the expression of EZH1, p65, and Pgf in hyperoxia + HDAC3 ^−/−^ + miR-17-antagomir-treated mice was similar to the hyperoxia-induced BPD mice (Fig. 6F). The above-mentioned results demonstrated that HDAC3 could regulate Pgf expression through the miR-17-EZH1-p65 axis to participate in alveolarization and pulmonary angiogenesis of BPD mice.

**Fig. 6.**
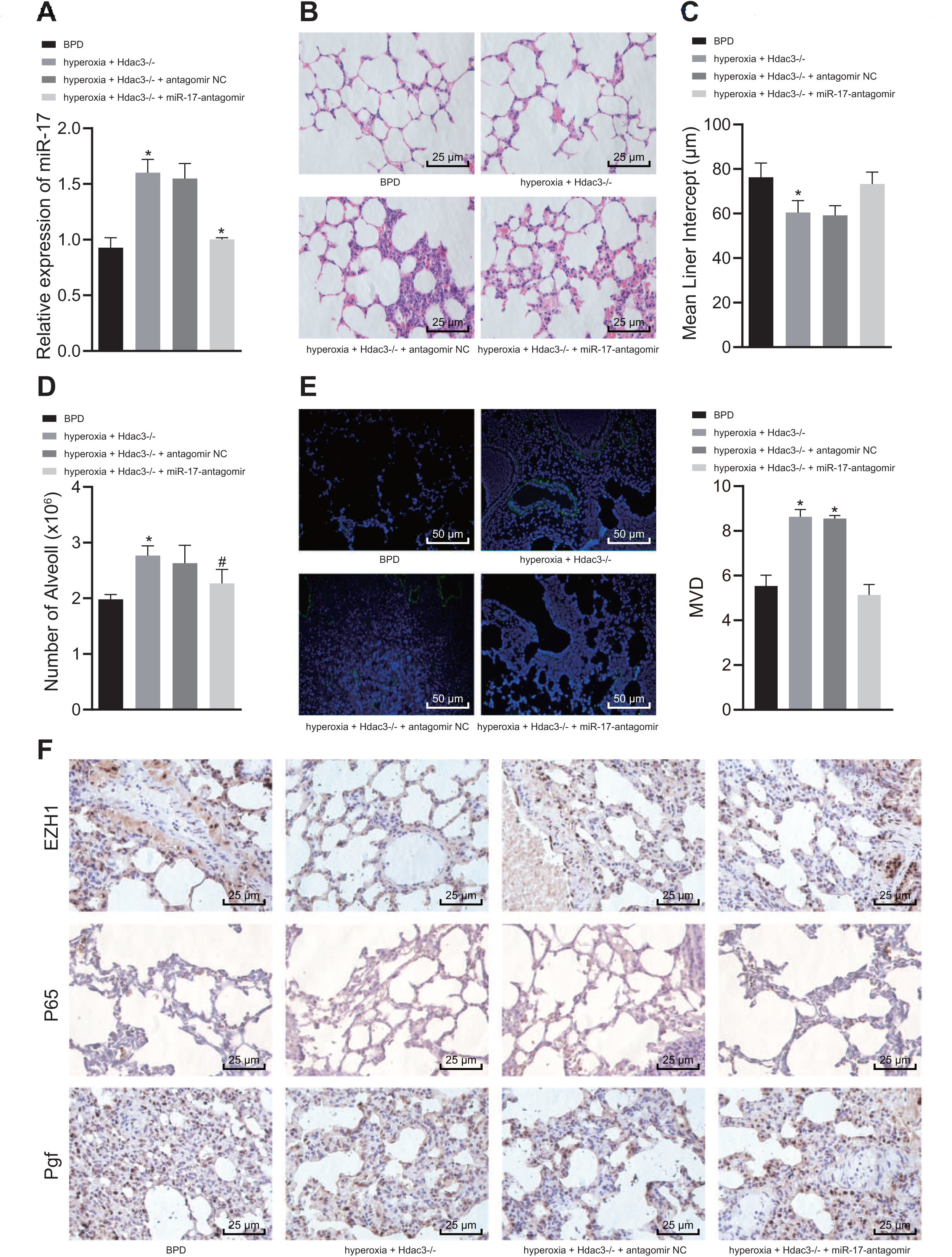
HDAC3 regulates Pgf expression through miR-17 to promote the alveolarization and pulmonary angiogenesis of BPD. A, The expression of miR-17 in different groups of mice detected by RT-qPCR. B, The degree of lung injury in mice of different groups detected by HE staining (× 400). C: The MLI of alveolar in the lung tissue of mice in each group. D, The changes in alveolar number in lung tissue of mice in each group. E, Immunofluorescence assay of MVD in lung tissues of mice in each group (× 200). F, The expression of EZH1, p65, and Pgf in lung tissues of mice following different treatment detected by immunohistochemistry (× 400). * *p* < 0.05. The above data were measurement data and expressed as mean ± standard deviation. Data among multiple groups were analyzed by one-way ANOVA, followed by Tukey’s post-hoc test. n = 10 for each mouse group.

## Discussion

Bronchopulmonary dysplasia (BPD) is a serious complication afflicting preterm infants which arises from oxygen toxicity and mechanical injury during oxygen supplementation [18]. Despite significant advances in the treatment of BPD, the mortality and morbidity of patients with BPD still very high, and yet the underlying mechanism of BPD progression remains poorly understood and inadequately defined [19]. Therefore, the discovery of therapeutic targets is critically needed to facilitate the diagnosis and treatment of patients with BPD. Existing literature has highlighted the participation of HDAC in various pathological and physiological processes including BPD [5]. In this study, we established *in vivo* BPD mouse models and identified that HDAC3 might be an endogenous regulator of BPD, as HDAC3 could promote the angiogenesis and alveolarization in BPD by enhancing Pgf *via* the miR-17-EZH1-p65 axis.

Initial results revealed that the expression of HDAC3 was highly expressed while the miR-17-92 cluster was poorly expressed in BPD mice. As one of the members in the class I histone deacetylase family, HDAC3 possesses regulatory function on gene expression through the deacetylation of both histones and non-histone proteins [20]. HDAC3 has been shown to be upregulated in adenocarcinoma of the lung, which was related to its poor prognosis [6]. In agreement with our findings, the expression of miR-17 in the miR-17-92 cluster has been found to be poorly expressed in infants diagnosed with severe BPD, and also was correlated with the diagnosis of BPD [21]. Previous evidence has shown that HDAC3 could participate in regulating the remodeling and expansion of distant alveolar vesicles into primitive lung alveologenesis through mediation of miR-17-92 cluster [14], suggesting that HDAC3 may exert a regulatory role in the angiogenesis and alveolarization of BPD through inhibiting the expression of miR-17-92 cluster.

Next, the results obtained from dual-luciferase reporter gene assay indicated that miR-17 could bind to the 3’-UTR of EZH1, demonstrating that EZH1 was a target gene of miR-17 and could be negatively regulated by miR-17. EZH1, a component of polycomb repressive complex 2, has been demonstrated to exert a significant role in repressing the transcription of target genes that affect the pathogenesis of various diseases [22]. The interaction between miRNA and its target mRNAs and their functional mechanisms in the pathophysiology of BPD have been addressed [23]. EZH1 has been previously reported to be a target gene of miR-17-5p, and the down-regulation of miR-17-5p could enhance the resistance to erlotinib in non-small cell lung cancer by regulating EZH1 [24], which was in consistent with our experimental results whereby miR-17 in the miR-17-92 cluster negatively regulated EZH1 expression, which reflects the biological characteristics of BPD.

Subsequently, we found out that EZH1 could promote Pgf expression by recruiting p65 and then participated in the regulation of BPD. As an important molecule in angiogenesis, Pgf/Plgf has a regulatory effect on pathological conditions including tumor formation, ischemia, cardiovascular diseases, suggesting that it may serve as a novel therapeutic target in various diseases [25]. Moreover, Pgf has an essential role in pulmonary vascular development of BPD [26]. According to previous literature, it was demonstrated that EZH1 could mediate the recruitment of p65 to promote the transcription of target genes [11]. In addition, p65 could directly bind to the Pgf promoter region, leading to a substantial activation of Pgf transcript levels [12]. These results were in line with our results and the interaction of EZH1, p65 and Pgf in regulating the angiogenesis and alveolarization of BPD were verified. Another important finding was that HDAC3 could stimulate Pgf expression *via* the miR-17-EZH1-p65 axis, thus promoting the progression of BPD.

## Conclusions

In conclusion, the key findings obtained from the current study was the identification of the regulatory role of HDAC3 interacting with the miR-17-EZH1-p65-Pgf axis in pulmonary angiogenesis and alveolarization in newborn mice with BPD (Fig. 7). The upregulation of HDAC3 and inhibition of miR-17 expression were observed in BPD mice. In addition, the binding of miR-17 to EZH1 could regulate the recruitment of p65 to enhance the transcription of target gene Pgf, thereby regulating BPD. These findings suggest that HDAC3 may serve as a promising diagnostic biomarker and therapeutic target for BPD. At present, the effects and mechanisms of HDAC3 interacting with miR-17-EZH1-p65-Pgf in BPD remain scantly identified. Therefore, more in-depth investigation is urgently needed to further discuss the underlying rules that govern their interaction, which might increase the feasibility and safety of its therapy in clinical applications.

**Fig. 7.**
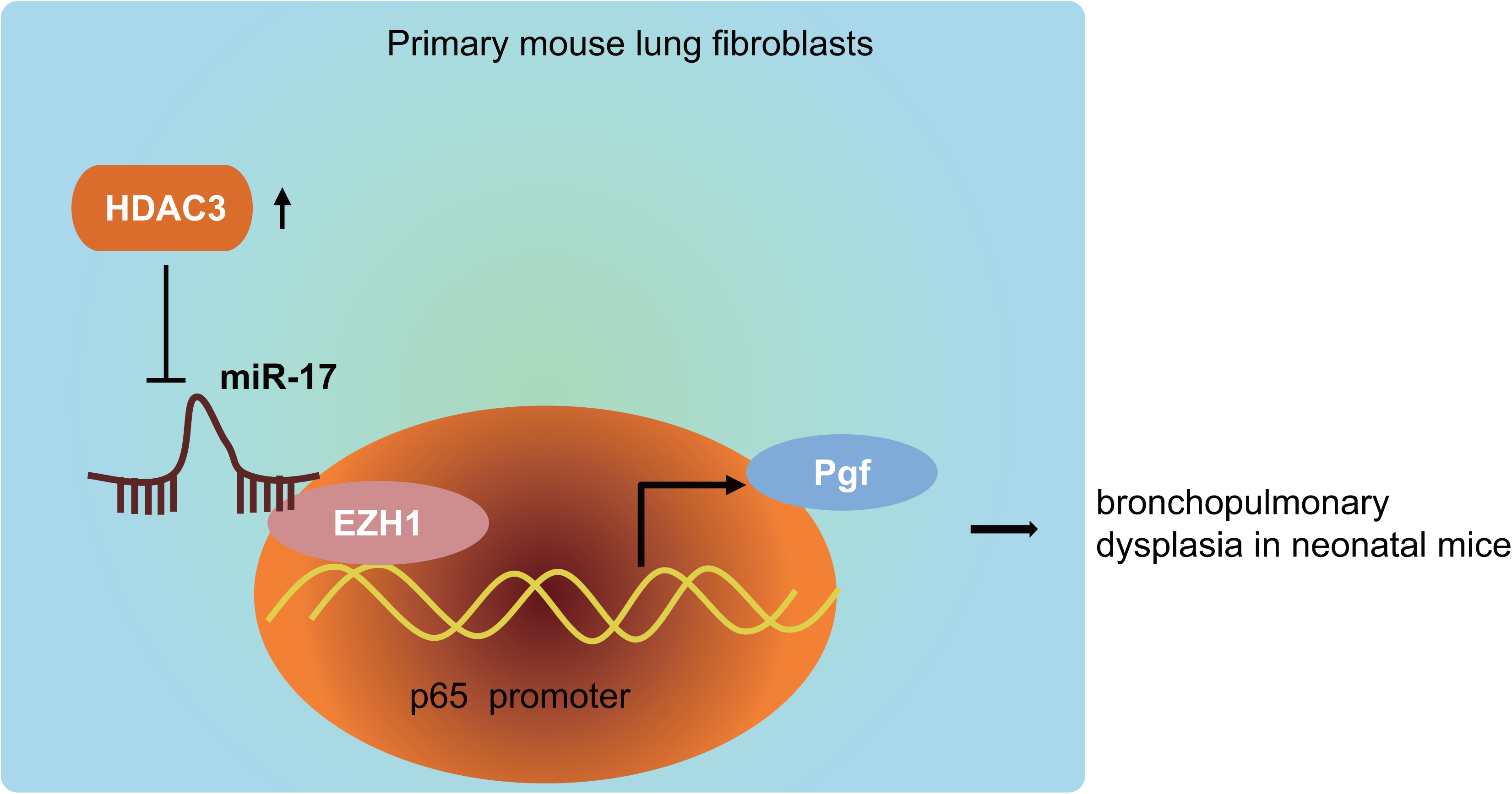
Schematic diagram depicting the role of HDAC3 regulating the miR-17-EZH1-p65-Pgf axis in BPD in neonatal mice. HDAC3 was highly expressed in lung fibroblasts of BPD mice, inhibiting the expression of miR-17 in the miR-17-92 cluster. miR-17 bound to EZH1 and regulated the transcription and expression of Pgf, a downstream target gene of transcription factor p65 and thereby regulating the development of BPD.

## Abbreviations

BPD: Bronchopulmonary dysplasia
HDACs: Histone deacetylases
miR: microRNA
FBS: fetal bovine serum
HE: Hematoxylin-eosin
SP: streptavidin-peroxidase
MVD: microvessel density
RT-qPCR: Reverse transcription-quantitative polymerase chain reaction
cDNA: complementary DNA
GAPDH: Glyceraldehyde-3-phosphate dehydrogenase
TBST: Tris-buffered saline Tween-20
SDS-PAGE: sodium dodecyl sulfate-polyacrylamide gel electrophoresis
ChIP: Chromatin immunoprecipitation
assay NaCl: Sodium chloride
EDTA: ethylenediamine tetraacetic acid
PIC: pseudoisocyanine
IgG: immunoglobulin G
3’UTR: 3’untranslated region
ANOVA: analysis of variance

## Declarations

### Author contributions

DW, HH, and XXL designed the study, and were involved in data collection. DW and JL performed the statistical analysis and preparation of figures. HH and ZQZ drafted the paper. All authors read and approved the final manuscript.

### Availability of data and materials

Data generated and analyzed as part of this study are included in the manuscript or are available upon request from the corresponding author.

### Consent for publication

Not applicable

## Acknowledgement

The authors sincerely appreciate all members participated in this work.

